# Pneumolysin is responsible for differential gene expression and modifications in the epigenetic landscape of primary monocyte derived macrophages

**DOI:** 10.1101/2020.06.08.139980

**Authors:** J. Cole, A. Angyal, R. D. Emes, T.J. Mitchell, M.J. Dickman, D.H. Dockrell

**Affiliations:** Department of Infection, immunity & Cardiovascular Diseases, University of Sheffield; Sheffield Teaching Hospitals NHS FT; The Florey Institute, University of Sheffield; Department of Chemical & Biological Engineering, University of Sheffield; Advanced Data Analysis centre, University of Nottingham; School of Veterinary Medicine and Science. University of Nottingham; Institute of Microbiology and Infection, University of Birmingham; MRC Centre for Inflammation Research, University of Edinburgh

## Abstract

Epigenetic modifications regulate gene expression in the host response to a diverse range of pathogens. The extent and consequences of epigenetic modification during macrophage responses to *Streptococcus pneumoniae*, and the role of pneumolysin, a key *Streptococcus pneumoniae* virulence factor, in influencing these responses, are currently unknown. To investigate this, we infected human monocyte derived macrophages (MDMs) with *Streptococcus pneumoniae* and addressed whether pneumolysin altered the epigenetic landscape and the associated acute macrophage transcriptional response using a combined transcriptomic and proteomic approach. Transcriptomic analysis identified 503 genes that were differentially expressed in a pneumolysin-dependent manner in these samples. Pathway analysis highlighted the involvement of transcriptional responses to core innate responses to pneumococci including modules associated with metabolic pathways activated in response to infection, oxidative stress responses and NFκB, NOD-like receptor and TNF signalling pathways. Quantitative proteomic analysis confirmed pneumolysin-regulated protein expression, early after bacterial challenge, in representative transcriptional modules associated with innate immune responses. In parallel, quantitative mass spectrometry identified global changes in the relative abundance of histone post translational modifications (PTMs) upon pneumococcal challenge. We identified an increase in the relative abundance of H3K4me1, H4K16ac and a decrease in H3K9me2 and H3K79me2 in a PLY-dependent fashion. We confirmed that pneumolysin blunted early transcriptional responses involving TNF-α and IL-6 expression. Vorinostat, a histone deacetylase inhibitor, similarly downregulated TNF production, reprising the pattern observed with pneumolysin. In conclusion, widespread changes in the macrophage transcriptional response are regulated by pneumolysin and are associated with global changes in histone PTMs. Modulating histone PTMs can reverse pneumolysin-associated transcriptional changes influencing innate immune responses, suggesting that epigenetic modification by pneumolysin plays a role in dampening the innate responses to pneumococci.

**Author summary:** Pneumolysin is a toxin that contributes to how *Streptococcus pneumoniae*, the leading cause of pneumonia, causes disease. In this study, the toxin alters gene expression in immune cells called macrophages, one of the first lines of defence against bacteria at sites of infection. Modulation involved multiple immune responses, including generation of chemical signals coordinating responses in immune cells termed cytokines. In addition, changes were observed in histone proteins that are involved in controlling gene expression in the cell. Pneumolysin reduced early production of the cytokine TNF-α and a medicine vorinostat that modifies these ‘epigenetic’ histone modifications had a similar affect, suggesting epigenetic mechanisms contribute to the ability of pneumolysin to reduce immune responses.

## Introduction

Pneumolysin (PLY) is one of the key virulence factors of *S. pneumoniae* (the pneumococcus), the leading cause of community-acquired pneumonia (Kadioglu et al. 2008) and is present in the majority of clinical isolates causing invasive pneumococcal disease (IPD) (Gray, Converse, and Dillon 1980; D.-K. Hu et al. 2015). Pneumolysin is a cholesterol-dependent cytolysin and part of a family of toxins expressed in Gram-positive bacteria (Tweten 2005). It is a 53 kDa protein that contains four domains (Walker et al. 1987). One mechanism of action is through pore formation (van Pee et al. 2017) but increasingly it is recognized to mediate additional actions independent of the ability to form pores (Bewley et al. 2014).

In murine models of bacteraemia PLY sufficient mutants are associated with increased lethality compared to PLY deficient mutants, linking the toxin to virulence (Benton, Everson, and Briles 1995). Furthermore, the transmission of *S. pneumoniae* between hosts has been linked to the presence of inflammation in the nasopharynx and pneumolysin has been shown to promote inflammation, increase transmission and foster the survival *ex vivo* of *S. pneumoniae* (Zafar et al. 2017). It has been also suggested that pneumolysin facilitates blood stream invasion by *S. pneumoniae* (D.-Hu et al. 2015). This highlights its importance as a key virulence factor of *S. pneumoniae* due to its role in the transmission of *S. pneumoniae* between hosts, in the progression from nasopharyngeal colonisation to invasive pneumococcal disease, the stimulation of inflammation and its cytotoxic effects (Kadioglu et al. 2008).

Pneumolysin has been shown to alter a variety of immune responses. For example it has been demonstrated to both activate the classical complement pathway (Paton, Rowan-Kelly, and Ferrante 1984) as well as play a role in complement evasion (Quin, Moore, and McDaniel 2007). Pneumolysin has been associated with stimulation of NRLP3 and potentially other inflammasome components (McNeela et al. 2010; R. Fang et al. 2011). In human monocytes it is associated with production of tumour necrosis factor alpha (TNF-α) and interleukin (IL-)1β (Houldsworth, Andrew, and Mitchell 1994). Pneumolysin has been shown to be responsible for the differential expression of multiple genes in undifferentiated THP-1 cells, a monocytic cell line, (Rogers et al. 2003) but its impact on gene expression in primary macrophages has not been established in detail. More recently, pneumolysin has been shown to bind the mannose receptor C type 1 in mouse alveolar macrophages leading to diminished pro-inflammatory cytokine release and enhanced bacterial survival (Subramanian et al. 2019).

The subversion of the host immune system is one of the key components of bacterial pathogenesis. Microorganisms can highjack host gene expression to their benefit (Eskandarian et al. 2013; Hamon et al. 2007; Rolando et al. 2013). Epigenetic mechanisms such as histone post-translational modifications (PTMs) have been shown to regulate gene transcription (Berger 2007; Jenuwein and Allis 2001). Moreover, they can be modulated by a number of different bacterial components (Cole et al. 2016). Experiments using both *Legionella pneumophila* (Rolando et al. 2013), and *Listeria monocytogenes* (Eskandarian et al. 2013) have shown that bacterial interaction with the THP-1 monocytic cell line also modifies histone PTMs. In the case of *Listeria monocytogenes*, Listeriolysin O, a pore-forming cytolysin similar to PLY, is secreted causing dephosphorylation at serine 10 of histone H3 and reduction in the levels of acetylated H4 in THP-1 cells. In HeLa cells these changes altered the transcriptional profile, which was associated with a decrease in IL-6 and other genes involved in innate immune responses (Hamon et al. 2007). This supports the hypothesis that bacteria alter the epigenetic profile of the host cell as a strategy for immune subversion, limiting the inflammatory response to increase their survival.

The consequences of epigenetic changes during acute bacterial infections are, however, not fully understood. The aim of this study was to investigate the consequences and mechanism underpinning the ability of pneumolysin to modulate translational responses to *S. pneumoniae* using primary human macrophages. We found that pneumolysin modulates a broad range of immune transcriptional responses in macrophages and differential protein expression analysis also confirmed changes in key transcriptional modules. Moreover, we identified global changes in the abundance of histone PTMs in MDMs in a pneumolysin dependent manner. We illustrate that one key pneumolysin-regulated immune response, early TNF-α production is also altered by chemical manipulation of histone PTMs.

## Results

### Pneumolysin shapes the macrophage transcriptome in response to pneumococci

Transcriptional responses to Gram-positive bacteria occur rapidly and are well developed after three hours of bacterial challenge (Hamon et al. 2007)(Rogers et al. 2003). Therefore, we initially analyzed transcriptional changes in monocyte-derived macrophages (MDMs) three hours after bacterial challenge with either wild-type or a pneumolysin deficient (ΔPLY) strains. We confirmed that intracellular viability of pneumococci was equivalent at this time between wild-type and ΔPLY strains (Fig S1), thereby excluding potential confounding by different levels of vita-pathogen-associated molecular pattern (Sander et al. 2011).

We identified 1872 probe-sets that were differentially expressed in response to bacterial challenge with either strain (F value <0.05) and 1553 after multiple test correction false discovery rate (FDR <0.05). Next, we calculated moderated t tests for each infection, as shown in the Volcano plots in Figure 1. The analysis identified 503 probe-sets which are differentially expressed in a pneumolysin-dependent manner and 234 in an independent manner (summarised Fig S2). To provide further insight into the pneumolysin-dependent differential gene expression in MDMs we undertook pathway analysis of the differentially expressed genes.

**Figure 1:**
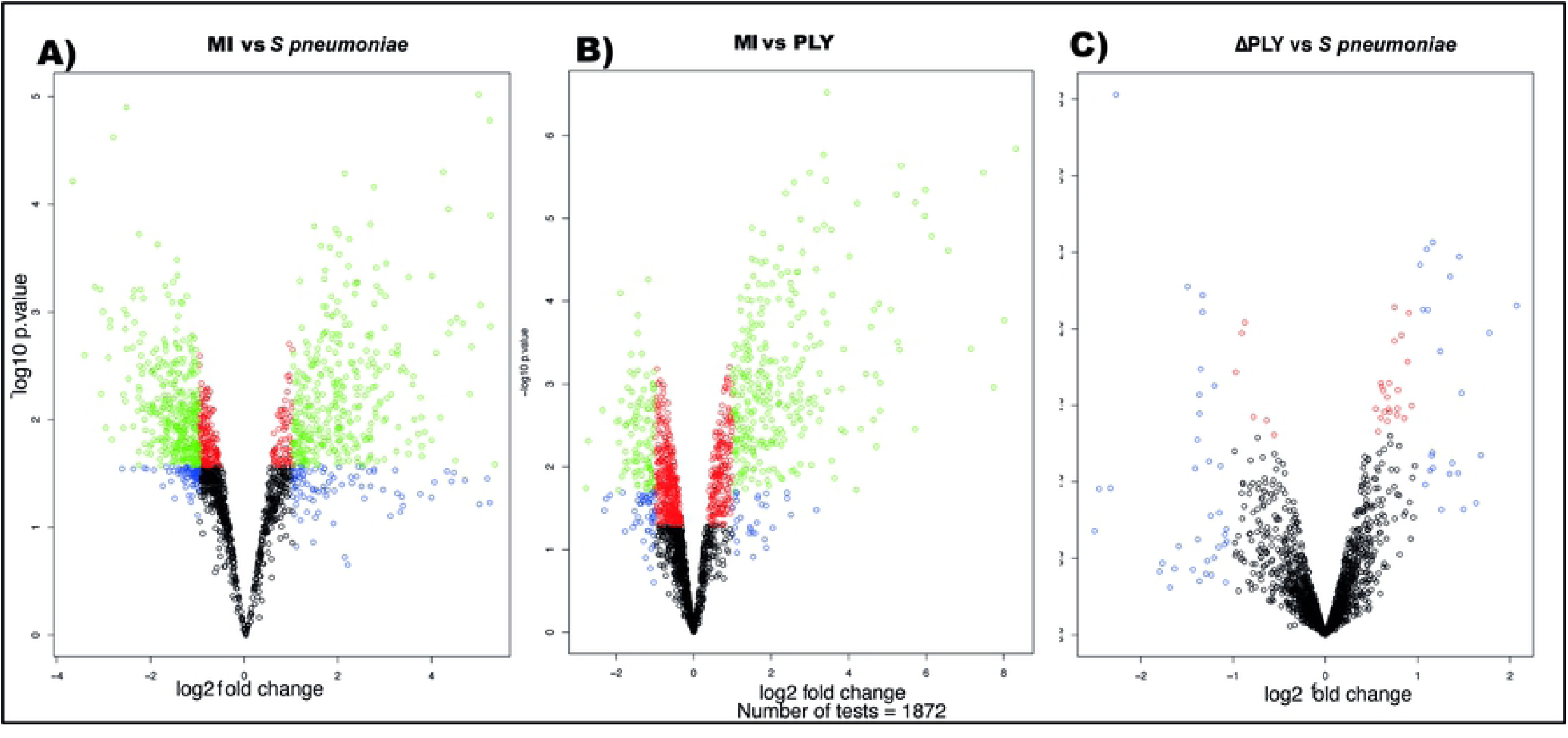
*S. pneumoniae* induced differential gene expression. Monocyte derived macrophages (MDMs) were challenged with either *S. pneumoniae*, the isogenic pneumolysin negative mutant (ΔPLY) or mock infected with phosphate buffered saline (MI) in biological replicates of three. After 3 h incubation the gene expression was measured using Affymetrix arrays. A) volcano plots comparing log2 (fold change) to log10 p value for MI vs *S. pneumoniae* challenged MDMs. B) volcano plots comparing log2 (fold change) to log10 p value for MI vs ΔPLY challenged MDMs. C) volcano plots comparing log2 (fold change) to log10 p value for *S. pneumoniae* vs ΔPLY challenged MDMs. The probe-sets in blue have a p value <0.05, in red an adjusted p value <0.05 (following FDR correction) and in green an adjusted p value <0.05 and an absolute fold change greater than 1.

### Pathway analysis of differentially expressed genes

Gene Ontology (GO) pathway analysis demonstrated that the top ten enriched terms in the pneumolysin-dependent differentially expressed genes were predominantly related to cell metabolism (Fig S3). However, there were also a number of terms relating to cellular responses to “stress” (17 in the *S. pneumoniae* challenge and 15 in the ΔPLY mutant) and in particular to oxidative stress responses. The volcano plots for the differentially expressed genes belonging to the GO term for oxidative stress response were plotted for each strain (Fig 2). These results highlighted that the differentially expressed genes were predominantly upregulated. Moreover, the TNF, heme oxygenase 1 (HMOX1) and prostaglandin-endoperoxide synthase 2 (PTGS2) genes were strongly upregulated.

**Figure 2:**
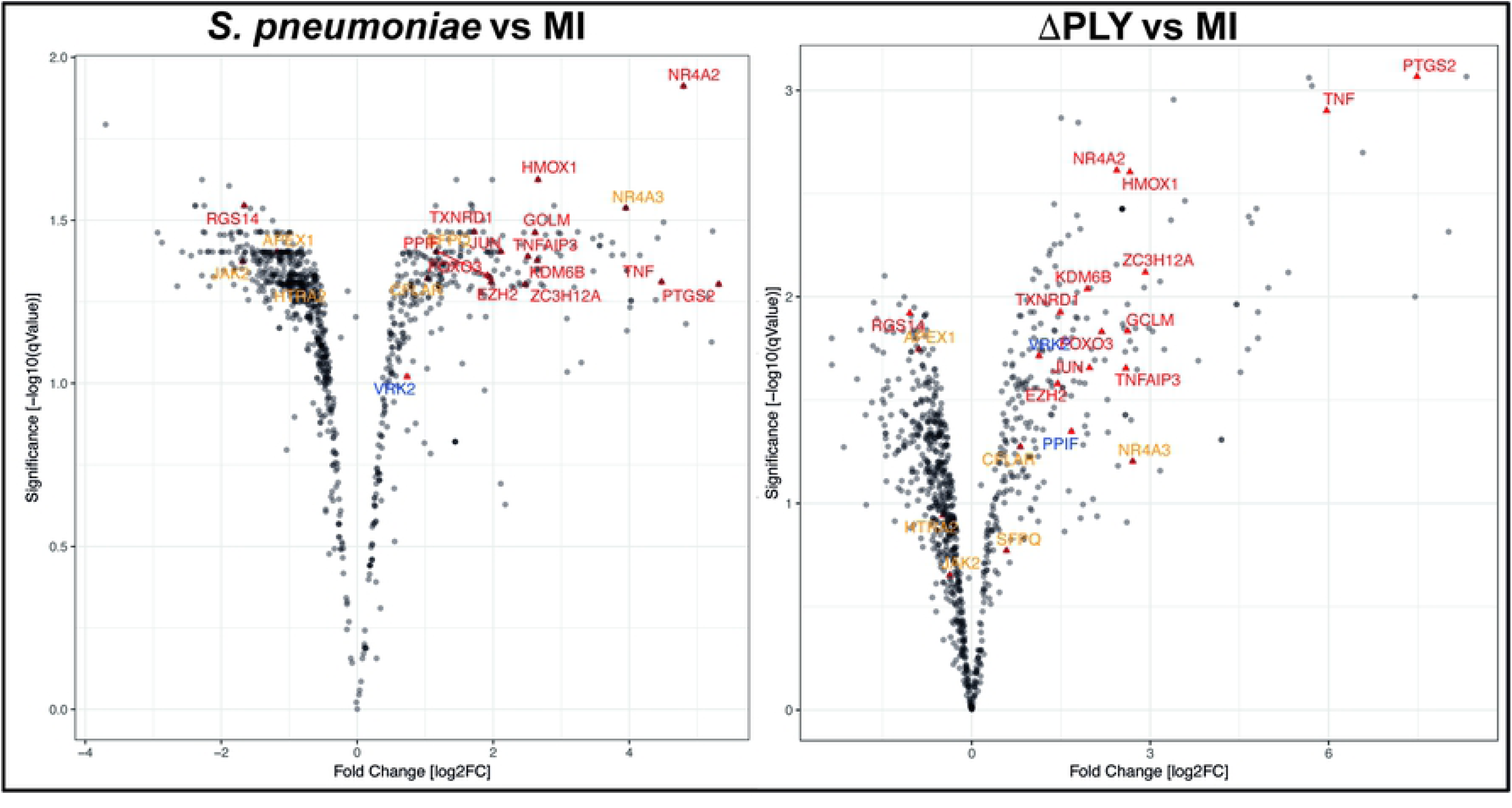
Oxidative stress response pathway terms significantly upregulated. Monocyte derived macrophages (MDMs) were challenged with either *S. pneumoniae*, the isogenic pneumolysin negative mutant (ΔPLY) or mock infected with phosphate buffered saline (MI) in biological replicates of three. After 3 h the gene expression was measured using Affymetrix arrays. A) volcano plots comparing log2 (fold change) to log10 p value for MI vs *S. pneumoniae* challenged MDMs. B) volcano plots comparing log2 (fold change) to log10 p value for MI vs ΔPLY challenged MDMs. The highlighted genes belong to the Oxidative stress response Gene Ontology term and are significantly expressed (q value <0.05). In red are those found in challenge with both strains, in blue the genes that are only found in the response to ΔPLY challenge and in orange the pneumolysin-dependent genes found after *S. pneumoniae* but not ΔPLY.

In addition to performing Gene ontology analysis we also analysed the differentially expressed genes using eXploring Genomic Relations (XGR) which performs hypergeometric enrichment analysis with background correction using the expression of monocyte derived macrophage cells to give a more cell type specific analysis. Canonical pathway analysis was performed for genes whose expression was upregulated in response to challenge with *S. pneumoniae* (adjusted p value <0.05). This revealed 36 over-represented pathways (Table S1). Importantly this demonstrated that both the TNF and the NFκ B signalling pathways were enriched. Analysis of the 2-fold downregulated terms did not reveal any enriched canonical pathways.

The analysis of the differentially expressed genes whose expression was up-regulated by greater than 2-fold in response to challenge with the ΔPLY mutant also revealed that the pathways for TNF, NFκB and CD40 signalling were significantly over-represented (Table S2). This highlights the importance of these pathways in the response to bacterial challenge. It also suggests that the increase in pro-inflammatory signals being released in particular TNF is independent of PLY. Of note, we observed a 6-fold increase in the TNF mRNA in response to ΔPLY but only 4-fold increase in response to the parent strain (Fig 2).

### Label-free quantitative proteomic analysis

Having established that there was a transcriptional difference in the host cell response following challenge with *S. pneumoniae* or ΔPLY, further studies were performed to examine the effects of *S. pneumoniae* on the proteome of MDMs.

Label free quantitative mass spectrometry was used to identify differentially expressed proteins following challenge with *S. pneumoniae* or ΔPLY. We identified 1807 proteins in samples challenged with *S. pneumoniae* and 1862 in those challenged with ΔPLY. On average we identified 1812 proteins in each of the 12 samples from 4 biological replicates that we analysed. Of these, we identified 32 that were differentially expressed between the three conditions with an F statistic value of less than 0.05 (summarised in Table 1). The results show that 16 proteins are differentially expressed in response to infections with *S pneumoniae*, and 22 in response to ΔPLY. Eight proteins were differentially expressed in a pneumolysin-dependent manner, these included serine / threonine protein kinase 1, also known as the oxidative stress response 1 protein (OXSR1) and Ubiquitin domain-containing protein 1 / 2 (UBTD2), and 9 were common to both bacterial strains, including Galectin 3 binding protein which is involved in the host immune responses (Breuilh et al. 2007) and the 26S proteasome non-ATPase regulatory subunit 5 (PSMD5). Several of the differentially expressed proteins identified such as PSMD5, UBTD2 and Ubiquitin conjugation factor E4 A (UBE4A) are involved in protein turn-over involving a number of pathways known to be important in the regulation of host immune responses (H. Hu and Sun 2016).

**Table 1:**
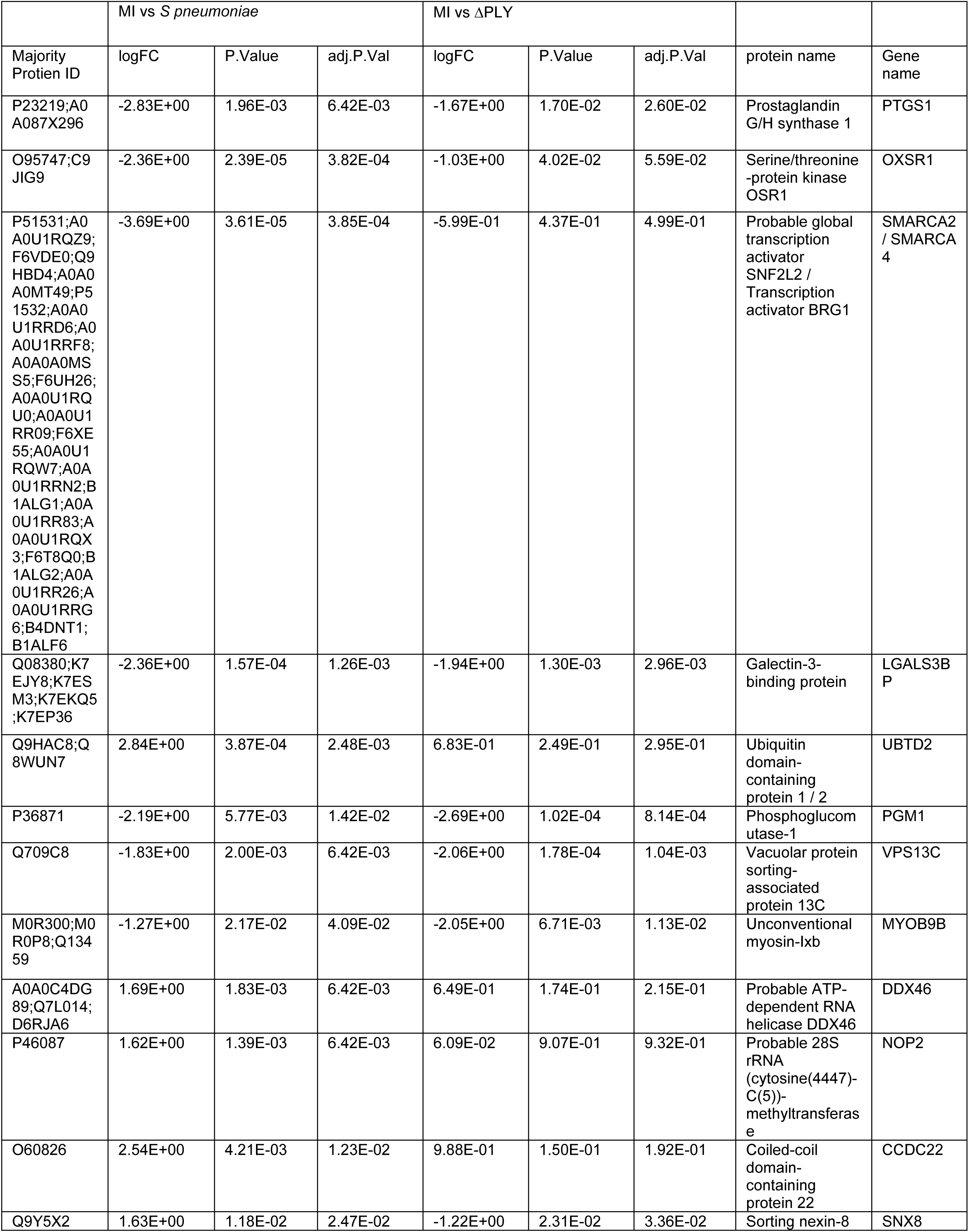
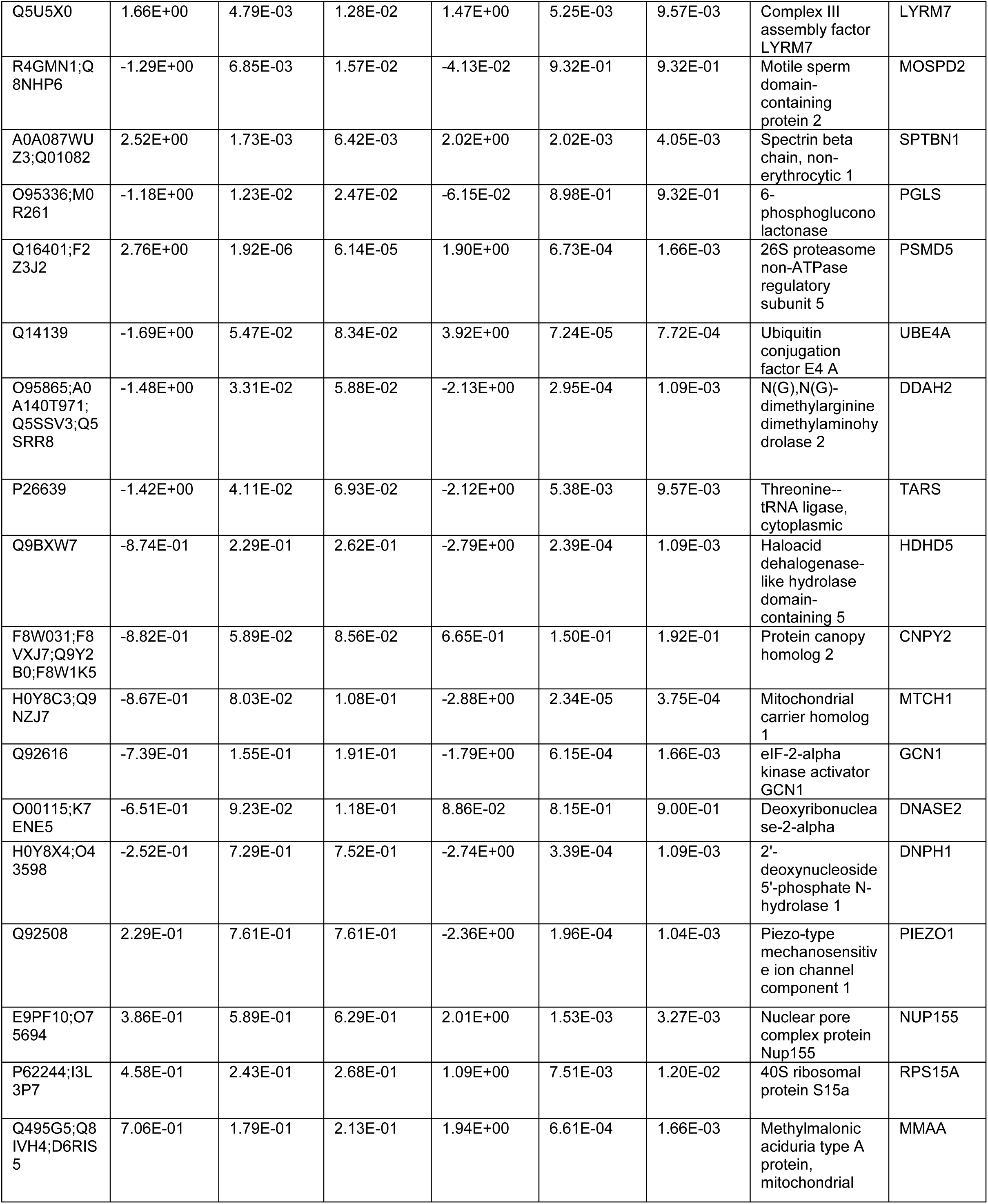
Differentially expressed proteins at 6 h following challenge with *S pneumoniae* or ΔPLY.

### Pneumolysin is responsible for changes in the relative abundance of histone post-translational modifications

Having defined a pneumolysin-dependent alteration of the macrophage transcriptome and proteome in response to *S. pneumoniae*, we sought next to establish if PLY also influenced epigenetic changes in response to *S. pneumoniae*. Quantitative mass spectrometry was used to identify changes in the global abundance of histone PTMs following challenge with *S. pneumoniae* or ΔPLY. Focussing on the most abundant PTMs, methylation and acetylation, on histones H3 and H4, in MDMs in response to bacterial challenge we identified 94 different peptide proteoforms and 18 were found to change in response to bacterial challenge (Fig 3 & 4). A subset of these changes were PLY-dependent, since they were significantly altered in *S. pneumoniae* in comparison to the ΔPLY mutant (Table 2). These included significant increases in the relative abundance of H3K4me1 and H3.3K36me2 following challenge with *S. pneumoniae* in comparison to ΔPLY. In addition, we observed decreases in the relative abundance of H3K9me2, H3K27me2 and H3K79me2 in a pneumolysin-dependent manner compared to both mock infected (MI; blue) and ΔPLY (Fig 3). Moreover, there was a pneumococcal associated PLY-independent increase in the level of H3K27me2K36me2 and a reciprocal drop in H3K27me3K36me1. We noted an increase in the relative abundance of H3.3K36me2 in a PLY dependent manner and an increase in the relative abundance of H3K23ac and an increase in H3.3K27me2K36me1 with a reciprocal drop in H3.3K27me2K36me2 in response to both bacterial challenges.

**Table 2:** Summary of the significant changes in relative abundance of histone PTMs identified following bacterial challenge

| Histone PTM | *S. pneumoniae* | ΔPLY |
| --- | --- | --- |
| H3K4me1 | Up | Down |
| H3K9me2 | Down | Up |
| H3K23ac | Up | Up |
| H3K27me2 | Down | Up |
| H3K27me2K36me2 |  | Up |
| H3K27me3K36me1 |  | Down |
| H3.3K27me2K36me1 | Up | Up |
| H3.3K27me2K36me2 | Down | Down |
| H3.3K36me2 | Up | Down |
| H3K79me2 | Down | Up |

**Fig 3:**
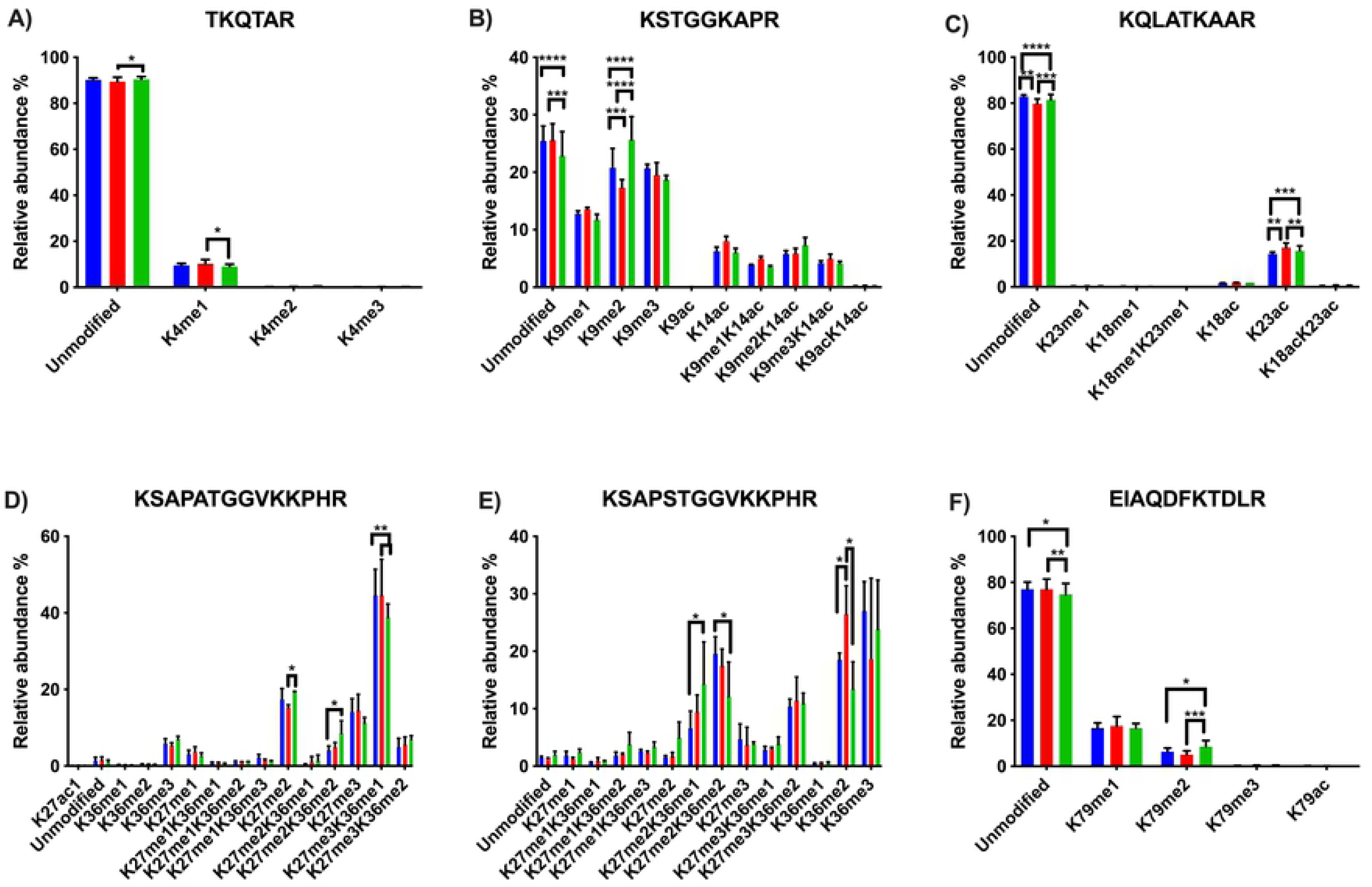
Pneumolysin is responsible for changes in relative abundance of PTMs on histone H3. Monocyte derived macrophages (MDMs) were challenged with either *S. pneumoniae*, the isogenic pneumolysin negative mutant (ΔPLY) or mock infected with phosphate buffered saline (MI) in biological replicates of three. After 3 h incubation the histones were extracted and analysed by mass spectrometry and the relative abundance of each post translational modification (PTM) was measured for each peptide. The bar plots represent the relative abundance of each PTM quantified, blue represents the abundance in MI MDMs, green following challenge with *S. pneumoniae* and in red to ΔPLY. A) Peptide TKQTAR shows an increase in the level of H3K4me1. B) Peptide KSTGGKAPR shows a decrease in the level of H3K9me2 in a pneumolysin-dependent manner. C) Peptide KQLATKAAR shows a relative increase in the level of H3K23ac in response to both bacterial challenges. D) Peptide KSAPATGGVKKPHR shows a decrease in H3K27me2 following challenge with *S. pneumoniae* compared to MI, an increase in the level of H3K27me2K36me2 and a reciprocal drop in H3K27me3K36me1 in response to ΔPLY challenge compared to MI. E) Peptide EIAQDFKTDLR shows an increase in the relative abundance of H3K79me2 following challenge with ΔPLY. F) Peptide KSAPSTGGVKKPHR of H3.3 shows an increase in H3.3K36me2 and in H3.3K27me2K36me1 with a reciprocal drop in H3.3K27me2K36me2. (One-way ANOVA, * p<0.05, ** p<0.01, *** p<0.001, error bars represent mean and standard deviation).

**Fig 4:**
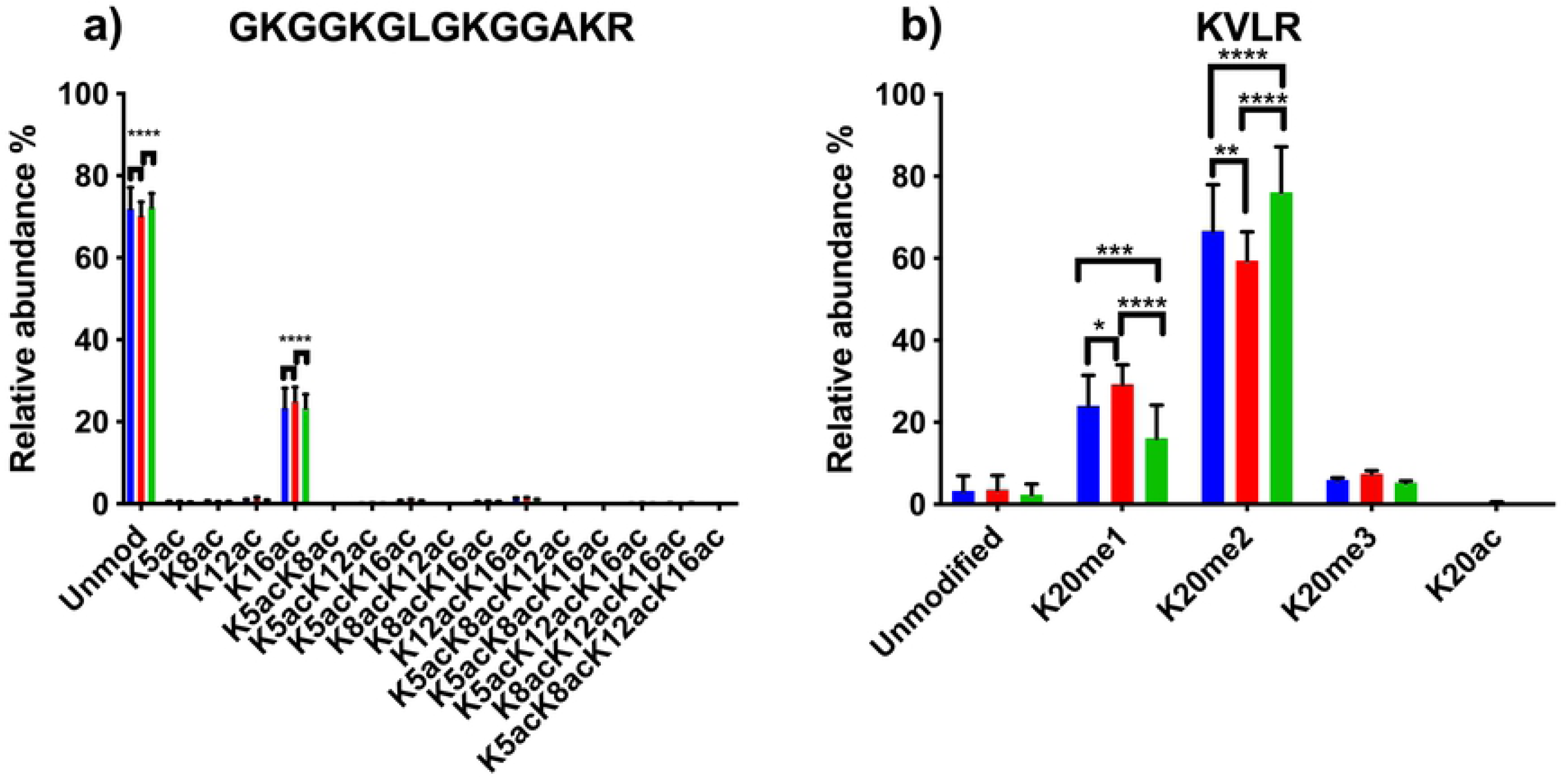
Pneumolysin is responsible for changes in relative abundance of PTMs on histone H4. Monocyte derived macrophages (MDMs) were challenged with either *S. pneumoniae*, the isogenic pneumolysin negative mutant (ΔPLY) or mock infected with phosphate buffered saline (MI) in biological replicates of three. After 3 h incubation the histones were extracted and analysed by mass spectrometry and the relative abundance of each post translational modification (PTM) was measured for each peptide. The bar plots represent the relative abundance of each PTM quantified, blue represents the abundance in MI MDMs, green following challenge with *S. pneumoniae* and in red to ΔPLY. A) Peptide GKGGKGLGKGGAKR shows a relative increase in the level of H4K16ac following challenge with *S. pneumoniae* in comparison to the ΔPLY. B) Peptide KVLR shows a decrease in the level of H4K20me2 in a pneumolysin-dependent manner compared and a reciprocal increase in the H4K20me1. Conversely for the MDM challenged with ΔPLY there was a rise in H4K20me2 and a fall in the level of H4K20me1.(n=3. One way ANOVA, * p<0.05, ** p<0.01, *** p<0.001, **** p<0.0001).

Further analysis of histone H4 showed a significant increase in the abundance of acetylation on K16 (H4K16ac) in response to infection with either strain (Fig 4). In addition, there was also a decrease in the dimethylated form of K20 (H4K20me2) and a reciprocal increase in the monomethylated form (H4K20me1) in a pneumolysin-dependent manner.

### Pneumolysin blunts inflammatory signalling during early infection

In order to establish the consequences of these changes to the transcriptome and epigenome, we next selected one prominent immune signature regulated by pneumolysin, TNF-α expression, and established this as a key component of the early immune response to *S. pneumoniae* (Jones et al. 2005). Using qRT-PCR we measured the abundance of TNFα mRNA in infected MDMs. There was a significant increase in the TNFα mRNA level following infection with the PLY deficient mutant but not the parent strain or the reconstituted mutant (Fig 5A). Next, we measured the amount of secreted of TNFα in MDM supernatants following 3 h of bacterial challenge (Fig 5B). We found that TNFα levels were significantly higher following challenge with the ΔPLY mutant suggesting that PLY initially blunts TNF-α expression.

**Fig 5:**
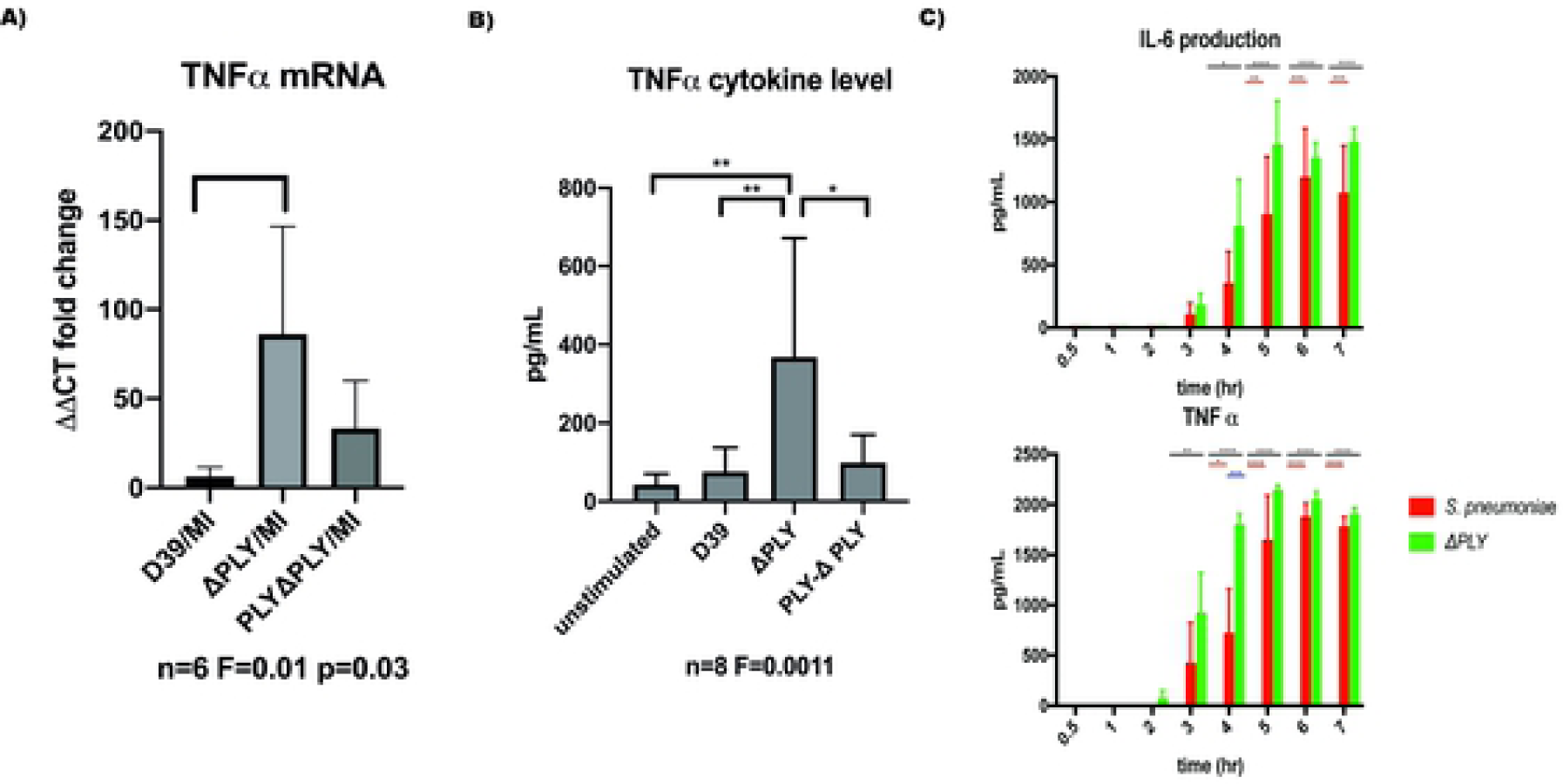
Pneumolysin leads to blunted TNFα release by repressing mRNA production. Monocyte derived macrophages (MDMs) were challenged with either *S. pneumoniae*, the isogenic pneumolysin negative mutant (ΔPLY), the isogenic pneumolysin negative mutant with reconstituted pneumolysin (PLY-ΔPLY). or mock-infected with phosphate buffered saline (MI). A) After 3 h incubation RNA was extracted from MDMs and RT-qPCR performed to measure the abundance of TNFα mRNA. The Bar chart represents the ΔΔCT fold change for each bacterial challenge demonstrating significantly higher abundance of TNFα mRNA following challenge with ΔPLY. One-way ANOVA F statistic =0.01 and p value <0.05. B) At 3 h TNFα concentration were measured in the supernatants of MDMs challenged with each strain. The Bar chart demonstrate the significantly raised level of TNFα in response to challenge with ΔPLY (n=8, one way ANOVA F statistic = 0.0011, p<0.05). C) TNFα and IL-6 cytokines were measured in supernatants following challenge with *S. pneumoniae* or ΔPLY at 0.5, 1, 2, 3, 4, 5, 6, 7 h. The bar charts represent mean and Standard Deviation of 3 biological replicates run in technical duplicates are shown demonstrating a significant difference in the rise of TNF-α between the two strains at 4 h, (black line highlights differences between control and ΔPLY, red line between control and *S pneumoniae*, blue line *S pneumoniae*, and ΔPLY, * p<0.05, ** p<0.01, **** p<0.0001).

Finally, we measured the concentration of TNFα over time using ELISA assays to determine if this initial anti-inflammatory effect was maintained (Fig 5C). The results showed that the difference seen in the release of TNF-α, was only observed at an early time window, 3-5 h after exposure to bacteria. This suggests that the anti-inflammatory effect of pneumolysin only plays a role over a defined time period.

### Histone deacetylase inhibitors reduce TNFα cytokines in response to infection

Having previously demonstrated that pneumolysin both modifies macrophage transcriptome and epigenome we next tested if epigenetic modifications could alter key PLY-associated immune responses. As a proof of concept, we pre-treated MDMs with the histone deacetylase inhibitor (HDACi) vorinostat (SAHA) or vehicle control (DMSO) following challenge with *Streptococcus pneumoniae* or ΔPLY mutant and quantified TNFα release (Fig 6). As previously shown, challenge with the ΔPLY mutant strain resulted in significantly increased TNFα levels as compared to the parental strain, but this level was significantly decreased in the ΔPLY mutant challenged cells by pre-treatment with vorinostat. This suggests that the epigenetic modifications we observed as being induced by PLY exposure could have functional consequences to immune responses, as illustrated for TNFα.

**Fig 6:**
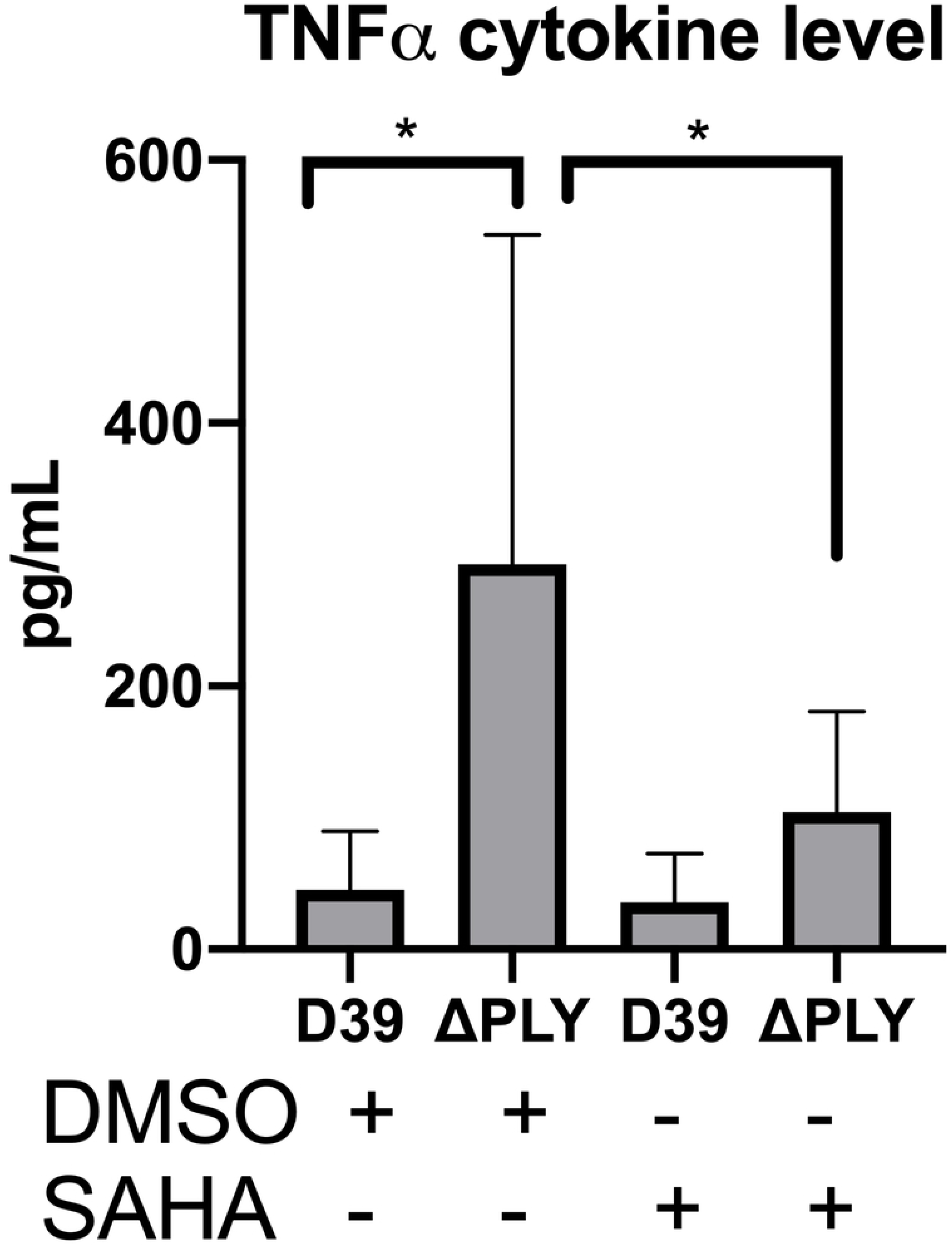
The HDAC inhibitor Vorinostat suppresses TNFα levels to similar levels as pneumolysin. Monocyte derived macrophages (MDMs) were challenged with either *S. pneumoniae*, the isogenic pneumolysin negative mutant (ΔPLY) or mock infected with phosphate buffered saline (MI) in the presence of 3 μM vorinostat (SAHA) or vehicle control (0.5% DMSO). The Bar chart and standard deviation represents the levels of TNFα in the supernatants demonstrating that suppression of TNFα can be mimicked by SAHA to similar levels as in the presence of PLY. (n=7 p<0.05).

## Discussion

Using a range of systems level approaches we have demonstrated that pneumolysin exerts a strong influence over the host response to *S. pneumoniae* at the transcriptomic, proteomic and epigenetic level. We have shown that 503 probes were differentially expressed in pneumolysin-dependent manner in association with global pneumolysin dependent changes in histone PTMs. These changes are associated with widespread changes in macrophage metabolism, oxidative stress responses and cytokine signalling and result in differential expression of key immune proteins, including TNF-α and IL-6. Crucially we show that use of a HDAC inhibitor to modify the epigenome is sufficient to reprise the reduction in TNF-α identified as occurring in response to pneumolysin.

Our results on the transcriptional response of macrophages to pneumolysin are consistent with previous transcriptomic analysis in undifferentiated THP-1 cells that showed 142 genes to be differentially expressed in a PLY dependent manner and 40 to be PLY independent (Rogers et al. 2003). Our study in primary cells, identifies many more genes differentially regulated enabling more in-depth analysis of changes at a pathway level. A number of differentially expressed genes are involved in the immune response such as TNF and HMOX1. Pathway analysis of the differentially expressed probes highlighted the importance of metabolic pathways in response to infection, and in particular the role for oxidative stress responses. The host’s oxidative stress responses have been highlighted as playing a key role in the host response to *S pneumoniae* in lung epithelial cells (Zahlten et al. 2015). Importantly, this module is upregulated in alveolar macrophages (AM) and plays a key role in preserving responses such as phagocytosis which may otherwise be altered by dysregulated oxidative stress in AM and MDM (Bewley et al. 2014; Belchamber et al. 2019). Furthermore, nuclear factor erythroid 2 (NRF2), the master regulator of antioxidant responses, plays a pivotal role in the protection against lung injury (Zhao et al. 2017), reduces lung inflammation after intratracheal instillation of LPS and decreases mortality in systemic models of inflammation (Thimmulappa et al., 2016). Crucially, NRF2-regulated pathways are amenable to therapeutic manipulation as evidenced by improved AM phagocytosis in patients with chronic obstructive pulmonary disease following *ex vivo* treatment with NRF2 agonists and also in a mouse model of cigarette smoke associated impairment of bacterial clearance in the lung (Harvey et al. 2011). The XGR pathway analysis following bacterial challenge further highlighted NFkB, NOD-like receptor and TNF signalling as enriched pathways emphasising their importance in the host’s response to bacteria.

The label-free quantitative proteomic analysis revealed a small number of significantly differentially expressed proteins, but substantiated involvement of several of the pathways identified at the transcriptomic pathway level. Examples of proteins linked to these pathways included an increase in the coiled-coil domain containing protein 22 (CCDC22), that plays an essential role in NFΚB signalling and whose depletion leads to blockade of signalling (Starokadomskyy et al. 2013). As an example of a protein downregulated we noted that the oxidative stress response protein 1 (OXSR1) involved in the oxidative stress response (W. Chen, Yazicioglu, and Cobb 2004) was decreased in a PLY-dependent manner.

The proteomic analysis also identified a number of differentially expressed proteins that are involved in regulating gene transcription and protein translation. The probable global transcription activator SNF2L2 / Brahma and the transcription activator Brahma-related gene 1 proteins were decreased in PLY dependent manner. They are associated with regulation of gene transcription by chromatin remodelling (B. G. Wilson and Roberts 2011). The ATP dependent RNA helicase DDX46 which is associated with pre-mRNA splicing (Will et al. 2002) was also increased in a PLY dependent manner. In addition, we observed an increase in the probable 28S rRNA (cytosine(4447)-C(5))-methyltransferase encoded for by the NOP2 gene which has been shown to be associated with the assembly of the large subunit of the ribosome and may play a role in cell cycle and proliferation (Sloan, Bohnsack, and Watkins 2013). In summary, the quantitative proteomic analysis identified a number of pneumolysin-dependent differentially expressed proteins involved in the regulation of gene transcription, protein translation, and modulation of signalling pathways such as the NFkB pathway, which are predicted to regulate innate immune responses to bacteria.

Although the absolute number of differentially expressed proteins is small, these results are in keeping with a number of previously published proteomic studies. In a study of THP-1 cells infected with *Mycobaterium tuberculosis* only 61 proteins were found to be differentially regulated (P. Li et al. 2017). In a study of alveolar macrophages infected with porcine reproductive and respiratory syndrome virus only 95 proteins were differentially expressed (Qu et al. 2017). Furthermore, this study has used primary cells and one of the reasons for the relatively small number of differentially expressed proteins may be the large differences between the biological replicates such that small differences in protein abundances following infection may not reach statistical significance. Several studies have shown only modest correlation between transcriptomic and proteomic datasets (Stare et al. 2017; G. Chen et al. 2002; Pascal et al. 2008; Ghazalpour et al. 2011), which might further explain the small number of differentially expressed proteins. Moreover, the transition from altered transcriptional response to altered proteome may take longer than the 6 hour time point studied. Indeed, in the study of proteomic response to *M. tuberculosis* and pathogenic porcine reproductive and respiratory syndrome virus both authors used a 24 hour time point (H. Li et al. 2017; Qu et al. 2017). It is also possible that many of the differentially expressed proteins are of relatively low abundance and therefore not readily detected using a shotgun proteomics approach.

Finally, the quantitative mass spectrometry analysis of histone PTMs in response to *S pneumoniae* was used to identify and quantify 94 different peptide proteoforms. Of these, there were 5 whose relative abundance changed in a pneumolysin dependent manner. This is to our knowledge the first time that this approach has been applied to study the changes in relative abundance of histone PTMs in primary MDMs in response to *S. pneumoniae*. The advantages of mass spectrometry analysis used in this study were emphasised by the identification of combinatorial marks on both H3 and H3.3 that would not have been possible using antibody-based approaches. Interestingly several of these modifications are associated with activating or repressing gene transcription. Although some histone PTMs are well characterised in a number of systems, their exact function in the context of infections is yet to be fully described. It is likely to vary between cell types. Furthermore, individual PTMs will not act in isolation but rather be part of a combinatorial code fine tuning responses to environmental stressors, such as to bacterial infection and at later time points after exposure to consequences such as DNA damage. Nevertheless, in light of the changes observed in response to challenge with *S. pneumoniae* (*e*.*g*. increases in H3K4me1, H3K23ac or H4K16ac and decreases in H3K9me2 or H3K79me2) it is possible that these changes represent the removal of repressive marks and the increase in marks associated with active gene transcription. This would allow fine tuning of the innate immune host response to bacteria to occur. This highlights the important role played by PTMs in the response to infection and therefore offer the potential for the novel use of therapeutic approaches involving immunomodulation of host responses through the use of emerging classes of drugs that target the enzymes that regulate these histone PTMs.

The functional consequences of the net changes to transcriptional responses involving inflammatory cell signalling and regulation of protein expression were examined focusing on TNF-α as an exemplar early response cytokine that plays a key role against *S. pneumoniae* (Jones et al. 2005). We selected this cytokine since PLY induced a significant temporal reduction in production of TNF-α, in conjunction with decreased production of IL-6. Our demonstration that pneumolysin inhibits the early production of TNF-α is consistent with the findings others have reported in dendritic cells and murine AM (Subramanian et al. 2019). We were able to demonstrate that changes in the epigenetic landscape are sufficient to reverse the maximal induction of TNF-α production observed following challenge with the PLY-deficient mutant. These experiments, using a HDACi, suggest that blunting of TNF-α release by MDM following challenge with a pneumococcal strain expressing PLY could be mediated through epigenetic mechanisms. This is important since early response cytokine generation is critical to the outcome of infection in *S. pneumoniae* and any modulation through reshaping the epigenetic landscape would be anticipated to have major consequences to the outcome of infection. Of note, the HDAC inhibitor Cambinol has been shown to inhibit TNF and IL-6 secretion in bone marrow-derived macrophages stimulated with LPS and was associated with greater survival in a murine lethal endotoxemia model and in response to *Klebsiella pneumoniae* challenge (Lugrin et al. 2013). In this case the therapeutic intervention was thought to limit excessive cytokine responses whereas in the case of PLY we would suggest it may limit early cytokine responses required to enhance early pathogen control. This emphasizes that in line with other aspects of host modulation through epigenetic manipulation would require careful calibration.

In summary our findings using a range of systems levels approaches show that pneumolysin modifies the early host response of macrophages to *S. pneumoniae* through modification of inflammatory responses, resistance to oxidative stress and metabolic responses. We observe that these transcriptional responses and the differential protein expression are associated with global changes in histone PTMs. We show alteration of the epigenetic landscape has functional consequences to key immune responses such as TNF-α production and suggest that PLY exerts these affects through epigenetic modulation and regulation of histone acetylation status. Our results also hint at a potential route by which these early host responses can be altered to improve responses to infection through the use of agents that modulate the enzymes that induce the signature pathogen-mediated epigenetic marks that adversely impact host responses.

## Materials and methods

### Bacterial Strains

*Streptococcus pneumoniae* serotype 2 strain D39 (D39), the isogenic pneumolysin-deficient mutant D39-ΔPLY (ΔPLY), which has a single amino acid substitution in the pneumolysin sequence generating a STOP codon and the reconstituted mutant PLY-D39-ΔPLY, which expresses PLY under an erythromycin promoter, were kindly obtained from Prof T. Mitchell (University of Birmingham). These were cultured and characterized as previously described (Bewley et al. 2014). Strains were grown in brain heart infusion broth and 20% FCS to mid exponential phase (with or without 1μg/mL erythromycin).

### MDM infection

Whole blood was obtained from healthy volunteers. Ethical approval was granted by South Sheffield Regional Ethics committee (07/Q2305/7). Peripheral blood mononuclear cells were separated by differential centrifugation using a Ficoll-Paque gradient and differentiated into monocyte derived macrophages (MDM) for 14 d as previously described in 24 well plates (Corning) (Collini et al. 2018). Bacteria were washed in PBS and re-suspended in RPMI 1640 supplemented with 10% pooled human immune serum (from previously vaccinated volunteers with demonstrable antibody levels to serotype 2 pneumococci) (Gordon et al. 2000). MDM were challenged with either opsonised D39, ΔPLY or PBS, at a MOI of 10, rested on ice for 1 h and incubated at 37°C in 5% CO_2_ for a further 3 h (Minshull et al. 2016). For certain experiments cells were treated with 3 μM vorinostat (SAHA, Sigma) or 0.5% DMSO (vehicle control) for 30 min prior to bacterial challenge and vorinostat reintroduced after bacterial challenge.

### *S. pneumoniae* internalisation assay

MDM were challenged with opsonized *S. pneumoniae* for 3 h then washed three times in PBS, incubated for 30 min in RPMI media (Lonza) with 40 units/mL of benzylpenicillin (Sigma) and 20 mg/mL gentamicin (Sanofi). The cells were then washed three times in PBS and incubated in 250 μL of 2% saponin (Sigma) for 12 min at 37°C in 5% CO_2_, then 750 μL of PBS was added, followed by vigorous pipetting. The number of internalised viable bacteria were measured by counting the number of colony forming units on Colombia blood agar (CBA) after 24 h incubation at 37°C in 5% CO_2_ contained in these lysates measured in triplicate.

### Cytokine measurements

Supernatants were obtained from MDM challenged with bacteria and were analysed as per the manufacturers protocol using either Tumour necrosis factor alpha (TNF-α (Ready-set-go!™, eBioscience)) or interleukin 6 (IL-6kit (Ready-set-go!™, eBioscience)). Briefly, 96 well ELISA plates were coated with 100 μL of 1x capture antibody overnight at 4°C, washed, and blocked in 200 μL assay diluent for 1 h at room temperature. After a wash, the supernatants were then added as were the standards (recombinant human TNF-α or recombinant human IL-6) and incubated for 2 h at room temperature. The wells were then washed and the detection antibody (biotin-conjugated anti-human TNF-α or anti-human IL-6) was added. After washing 100 μL avidin-horse radish peroxidase (HRP) was added to each well for 30 min at room temperature. Plates were then washed, prior to adding 100uL tetramethylbenzidine substrate solution for 15 at room temperature. The reaction was stopped by adding 2M sulphuric acid, the plate was then read at 450 nm using a Multiskan® EX plate reader (Thermo Scientific), and the data analysed in GraphPad Prism version 7.0c. (GraphPad Software).

### RNA Extraction

After 3 h of bacterial challenge, cells were washed and lysed in 600 μL Tri Reagent (Sigma) for 15min at room temperature before storing at −80 C. Ribonucleic acid (RNA) extraction was performed following the manufacturers guidelines for Direct-Zol™ RNA miniPrep (Zymo). Briefly, samples in Tri Reagent^®^ were centrifuged at 12 000*g* for 1 min, then the supernatant was transferred to a fresh 1.5 mL tube, 100% ethanol was added in a 1:1 ratio and well mixed. This was then transferred to the Zymo-spin™ column, centrifuged for 1 min, then washed using 400 μL Direct-Zol™ RNA pre-wash, centrifuged again at 12 000*g* for 1 min, then washed in 700 μL RNA wash buffer and again centrifuged at 12 000*g* for 1 min. Finally the RNA was eluted out in 50µl DNase/RNase free water.

### Microarray mRNA expression analysis

Affymetrix chip micro-array (Human Genome U133 plus 2.0 array, Santa Clara, CA) analysis of samples from three individuals was undertaken to characterise gene expression as previously described(Bewley et al. 2018). Data analysis was performed as previously described (Bewley et al. 2018). Briefly, CEL files were analysed using R version 3.4.0. CEL files were read using *Simpleaffy* version 2.52.0(C. L. Wilson and Miller 2005); background intensity correction, median correction and quantile probeset normalisation was performed using robust multi-array average expression with the help of probe sequence (GCRMA), using *AffyPLM* version 1.52.1(Bolstad et al. 2005). The quality control matrix was generated using *Simpleaffy* and *AffyPLM*. Principal component analysis was performed in R. The probes whose intensity were within the lowest 20^th^ centile were removed, using Dplyr version 0.7.1. Differential gene expression was calculated using *Limma* version 3.32.2.

### Pathway analysis

Pathway analysis was performed from the differentially-expressed gene lists generated. Hypergeometric tests were calculated in R, using *GO*.*db* version 3.4.1 to search the Gene Ontology (GO) database (Ashburner et al. 2000)for molecular function, cellular component, and biological process, using a p value cut off of 0.01 and a minimum of 3 genes. In addition, canonical pathway analysis was performed using XGR (1.1.5) (H. Fang et al. 2016) using all of the differentially expressed probes identified by ANOVA calculated in *Limma* as the test background.

### Real time quantitative polymerase chain reaction (RT-PCR)

The abundance of TNFα mRNA in bacterial exposed and mock-infected MDM was measured using qPCR. To perform cDNA synthesis a high-capacity cDNA reverse transcription kit (Applied Biosystems) was used to make complementary DNA for qPCR assay as per manufacturers protocol. The DNA products were quantified on QuantStudio5 (Abi) using GoTaq qPCR master mix (Promega). The reactions were prepared as per manufacturers protocol.

Primers used

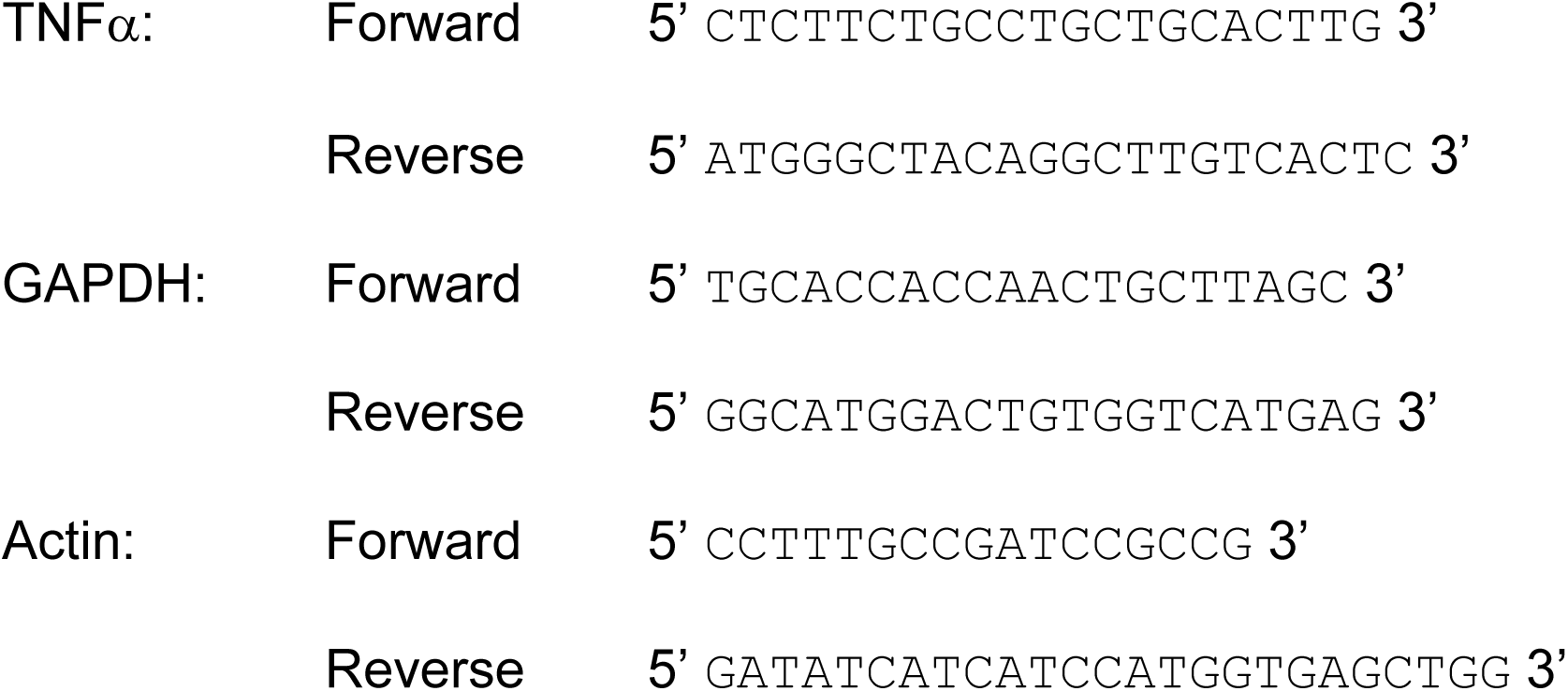

Fold change was calculated using Delta Delta Ct values.

### Quantitative proteomics

Protein lysates were digested with trypsin in conjunction with the Filter aided separation protocol (FASP) (Wiśniewski et al. 2009). Briefly, cells were lysed in 150 µL of 4% Sodium dodecyl sulphate (Sigma), 100 mM Tris HCl (Sigma) pH 7.6, 0.1 M Dithiothreitol (Sigma) and quantified using a BioRad DC assay as per the manufacturers protocol. 100 µg of protein lysates were mixed with 200 µL of 8 M urea dissolved in triethylammonium bicarbonate (TEAB) and added to the filter mounted in 11.5 mL low-bind Eppendorf. The tubes were centrifuged at 14 000 g for 30 min, the flow through discarded and further two 200 µL washes in 8M urea were performed. Then 100 µL 0.5 M iodoacetic acid was added for 20 min at room temperature and then centrifuged at 14 000 g for 30 min. The flow-through was discarded. The membrane was washed three times with 100 µL of 8 M urea, followed by three washes 100 mM TEAB. The lysates were trypsin digested overnight at 37 °C. The digested proteins were eluted in 120 µL 100mM TEAB. The samples were then desalted using Hypersep Hypercarb™ (ThermoScientific) tips following the manufacturers protocol. Briefly, tips were primed with elution solution (60% Acetonitrile 0.1% TFA) then washed in 0.1% TFA. The sample was then re-suspended on the tip by pipetting up and down 50 times. The tip was cleaned in 0.1%TFA, eluted 60% ACN 0.1% TFA and then 90% ACN 0.1% TFA. The samples were then dried down in Speedvac (Eppendorf). The peptides were re-suspended in 0.1%TFA and 3% ACN and loaded into and run on Ultimate 3000 RSLC nano flow liquid chromatography system with a PepMap300 C18 trapping column (Thermo Fisher), coupled to Q-Exactive HF Orbitrap mass spectrometer (Thermo Fisher). Peptides were eluted onto a 50 cm x 75 μm Easy-spray PepMap C18 column with a flow rate of 300 nL/min as previously described (Cole et al. 2019). Peptides were eluted using a gradient of 3% to 35% solvent B over 75 min. Solvents were composed of 0.1% formic acid (FA) and either 3% acetonitrile (ACN) (Solvent A) or 80% ACN (Solvent B). The loading solvent was 0.1% TFA and 3% ACN. Data acquisition was performed in full scan positive mode, scanning 375 to 1500m/z, with an MS1 resolution of 120 000, and AGC target of 1×10^6^. The top 10 most intense ions from MS1 scan were selected for Collision induced dissociation. MS2 resolution was of 30 000 with AGC target of 1×10^5^ and maximum fill time of 60 ms, with an isolation window of 2 m/z and scan range of 200-2000 m/z and normalised collision energy of 27.

### MaxQuant data analysis

The raw data from the MS were analysed in MaxQuant (version 1.5.6.5). The database search engine Andromeda was used to search the spectra against the UniProt database. The search settings were as follows: trypsin/P digestion, with up to 2 miss.ed cleavages, fixed modification was carbamidomethyl (C), variable modifications were oxidation (M) and acetylation (Protein N-term), Label Free Quantification (LFQ) was performed with a minimum number of neighbours of 3 and average number of neighbour of 6. Peptide tolerance was set at 4.5 ppm and minimum peptide length was set at 7, amino acid maximum peptide mass was set at of 4600 Da and Protein FDR was set at 0.01. Downstream analysis was performed in R version 3.4.0. The protein identification files were read in, results matching to a reverse sequence database and or those matching to a contaminant database were removed as were those with less than 2 unique peptides. The label-free intensities were then median-corrected for each sample and log2 transformed. Differential protein expression was calculated using Limma version 3.32.2. a repeated measures ANOVA to enable comparison to the microarray analysis (Goeminne, Gevaert, and Clement 2016).

### Histone extraction

Histone extraction was performed as previously described (Shechter et al. 2007). Briefly, MDMs were washed in PBS and scrapped in ice cold PBS with 1x protease inhibitors (Roche Complete EDTA free), before being pelleted at 900*g* for 10 min. Cell pellets were lysed in hypotonic lysis solution then re-suspended in 400 µL 0.2 M H_2_SO_4._ The histones were precipitated out by adding 132uL of 6.1N TCA to the supernatant and washed in acetone then resuspended in 100 μL of water (HPLC grade). Samples then underwent chemical derivitisation as previously described (Garcia et al. 2007). 10 μL of 100 mM ammonium bicarbonate pH 8 and 4 μl of ammonium hydroxide was added to 10 μg of histone sample. Then 10 μL of propionic anhydride in isopropanol (1:3 ratio) was added and 100% ammonium hydroxide used to keep the pH >8.0. The sample was incubated at 37°C for 15 min. Then it was dried down in a vacuum centrifuge (Concentrator plus, Eppendorf) and the process repeated. The samples were re-suspended in 40 μL of 100 mM ammonium bicarbonate and then tryptically digested overnight. The digestion was stopped by addition of glacial acetic acid and freezing at −80°C for 5 min. Finally, the samples were dried down before undergoing a further two rounds of proprionylation. Hypersep™ Hypercarb™ tip were used to desalt the samples of the chemical derivitisation residues following the manufacturers protocol for Hypersep ™ Hypercarb^™^ (ThermoScientific). (Minshull et al. 2016).

### Quantitative MS analysis of histone PTMs

The histone samples were re-suspended in 0.1% trifluoroacetic acid (TFA) and were analyzed on an Ultimate 3000 online nano-LC system with a PepMap300 C18 trapping column (ThermoFisher Scientific), coupled to a Q Exactive HF Orbitrap (ThermoFisher Scientific). Peptides were eluted onto a 50 cm × 75 μm Easy-spray PepMap C18 analytical column at 35°C. Peptides were eluted at a flow rate of 300 nL/min using a gradient of 3% to 25% over 55 min then 25% to 60% until 81 min. Solvents were composed of 0.1% formic acid (FA) and either 3% acetonitrile (ACN) (solvent A) or 80% ACN (solvent B). The loading solvent was 0.1% TFA and 3% ACN run in Data independent acquisition as previously described (Cole et al. 2019). The PTM identification and relative abundance was performed in Skyline and Epiprofile 2.0 also as previously described (Cole et al. 2019).

### Statistical analysis

Statistical analysis was performed using either R for the microarray and proteomic analysis, or for all other experiments in Prism version 7.0c (Graphpad). Data is presented as standard deviation (SD) or standard error of the mean (SEM). For all experiments a minimum of 3 biological replicates was used. Comparison between two paired groups employed a paired t-test, for comparison of 3 or more conditions a one-way analysis of variance (ANOVA) with Tukey’s post-test was performed. For comparison of multiple observations in more than two groups a two-way ANOVA with Tukey’s multiple comparison post-test was performed.

## Supplementary Figure legends

**Fig S1: Viable intracellular bacteria following 3 h challenge with *S. pneumoniae*, ΔPLY or PLY-ΔPLY mutants**.

Monocyte derived macrophages (MDMs) were challenged with either *S. pneumoniae*, the isogenic pneumolysin negative mutant (ΔPLY) or the isogenic pneumolysin negative mutant with reconstituted pneumolysin (PLY-ΔPLY). At 3 h the number of viable intracellular bacteria was determined. The results are expressed as mean and log_10_ of cfu/mL and represented as bar chart with standard deviation. (n=3, one way ANOVA p=0.99)

**Fig S2: Heatmap of the differentially expressed probes following MDM challenge**.

Monocyte derived macrophages (MDMs) were challenged with either *S. pneumoniae*, the isogenic pneumolysin negative mutant (ΔPLY) or mock-infected with phosphate buffered saline (MI) in biological replicates of three. At 3 h the gene expression was measured using Affymetrix arrays. This heatmap represents genes found to be differentially expressed (one-way ANOVA, with F p value <0.05).

**Fig S3: Gene ontology biological processes enriched terms show stress responses and metabolism are over-represented**.

Monocyte derived macrophages (MDMs) were challenged with either *S. pneumoniae*, the isogenic pneumolysin negative mutant (ΔPLY) or mock-infected with phosphate buffered saline (MI) in biological replicates of three. At 3 h the gene expression was measured using Affymetrix arrays. The bubble plot shows the top ten enriched Gene Ontology (GO) biological processes terms in both the pneumolysin mutant and the parental strain analysis. The bubble size and colour correspond to the number of genes which have mapped to the GO term. The bubbles are plotted along the x axis according to the –log10 p value for the enrichment.

**Table S1: Canonical pathway analysis of up-regulated differentially expressed probes following challenge of MDMs with *Streptococcus pneumoniae* using XGR**.

**Table S2: Canonical pathway analysis of up-regulated differentially expressed probes following challenge of MDMs with ΔPLY using XGR**

